# Fluorescent pH-sensitive nanosensors enable precise low-volume monitoring in high-throughput bioprocess manufacturing

**DOI:** 10.1101/2024.09.13.612862

**Authors:** Greta Csalane Besenyei, Alison Tang, Jonathan W Aylott, Veeren M Chauhan

## Abstract

Biopharmaceutical manufacturing requires precise pH control during production to ensure product quality and process performance. However, achieving this precision in low-volume, high-throughput automated settings present significant challenges with existing technology, often leading to inefficiencies and inaccuracies. This study introduces fluorescent nanosensors as a novel solution for accurate pH monitoring in micro-scale environments. Employing ratiometric fluorescence measurement, these nanosensors use analyte responsive and reference fluorophores in inert polyacrylamide matrices to perform dynamic measurements over pH 3.5 - 7.5. Nanosensors (35.26 ± 3.66 nm diameter) were synthesised with a neutral or positive surface charge (−5.08 ± 4.05 mV and +12.87 ± 1.25 mV, respectively). An automated workflow, using sacrificial and at line sample analysis, was developed by integrating the nanosensors with a TECAN™ automated liquid handling platform and a fluorescence spectrophotometer. The method was verified using in-process samples obtained from a monoclonal antibody purification, to highlight the compatibility of the nanosensors to different buffer systems in a typical biopharmaceutical manufacturing process. We show that pH-sensitive nanosensors can effectively monitor pH through the various stages of purification, demonstrating high accuracy (pH ± 0.23-0.40) even in sample volumes as low as 12.5 μL. The application of nanosensors represents a significant advancement in high-throughput scale-down bioprocess development by enabling precise, automated pH adjustment. This study improves the understanding of biopharmaceutical manufacturing, through the application of fluorescent nanosensors, paving-the-way for optimisations in low-volume and high-throughput product production.

**Graphical Abstract:** 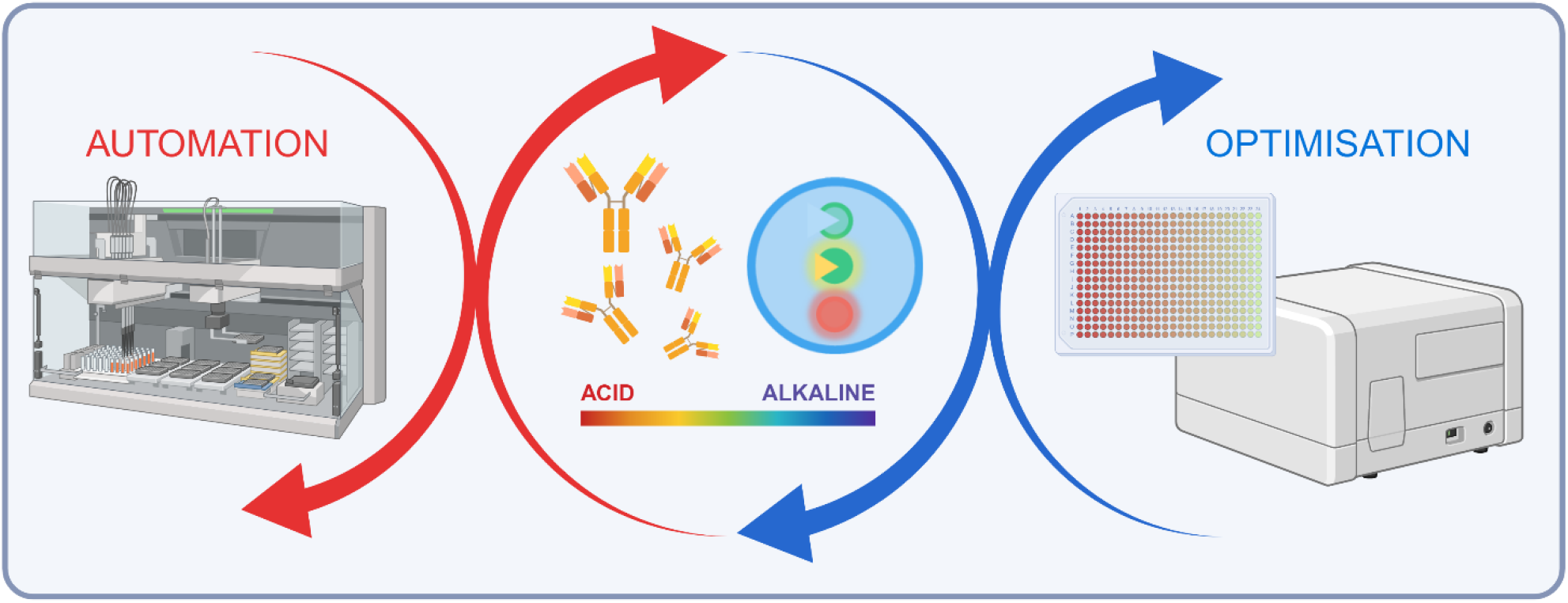

## Introduction

High-throughput process development (HTPD) is important for optimising bioprocesses, delivering substantial benefits in efficiency, cost savings, product quality, and innovation. Within HTPD, chromatography emerges as a focal point [1-4], serving as the key purification technique essential for eliminating process-related impurities (**Fig. 1a**). This highlights its critical role as the primary workhorse in the purification process, demonstrating its indispensable value in enhancing the purity and integrity of bioproducts. Efforts to expand HTPD to encompass viral filtration have increased [5], yet integrating individual unit operations into a fully automated workflow remains largely unexplored. A significant obstacle in achieving end-to-end bioprocess automation lies in the pH adjustment required between unit operations and for pH-based viral inactivation [6]. In addition, the challenge of measuring pH and acquiring real-time titration curve data in a high-throughput setting persists. While commercial solutions for non-invasive pH measurement exist, their accuracy and sensitivity fall short for bioprocessing applications [7]. Although high-throughput physical pH probes are available, their adoption is hampered by high costs and substantial capital investments needed for integration into automated liquid handling systems [8]. Developing a method for rapid, accurate pH measurement in a high-throughput context is important, particularly for scale-down purification processes reliant on automated systems. This advancement will be essential for streamlining process development workflows, thus enhancing the efficiency of bioprocesses.

**Fig. 1.**
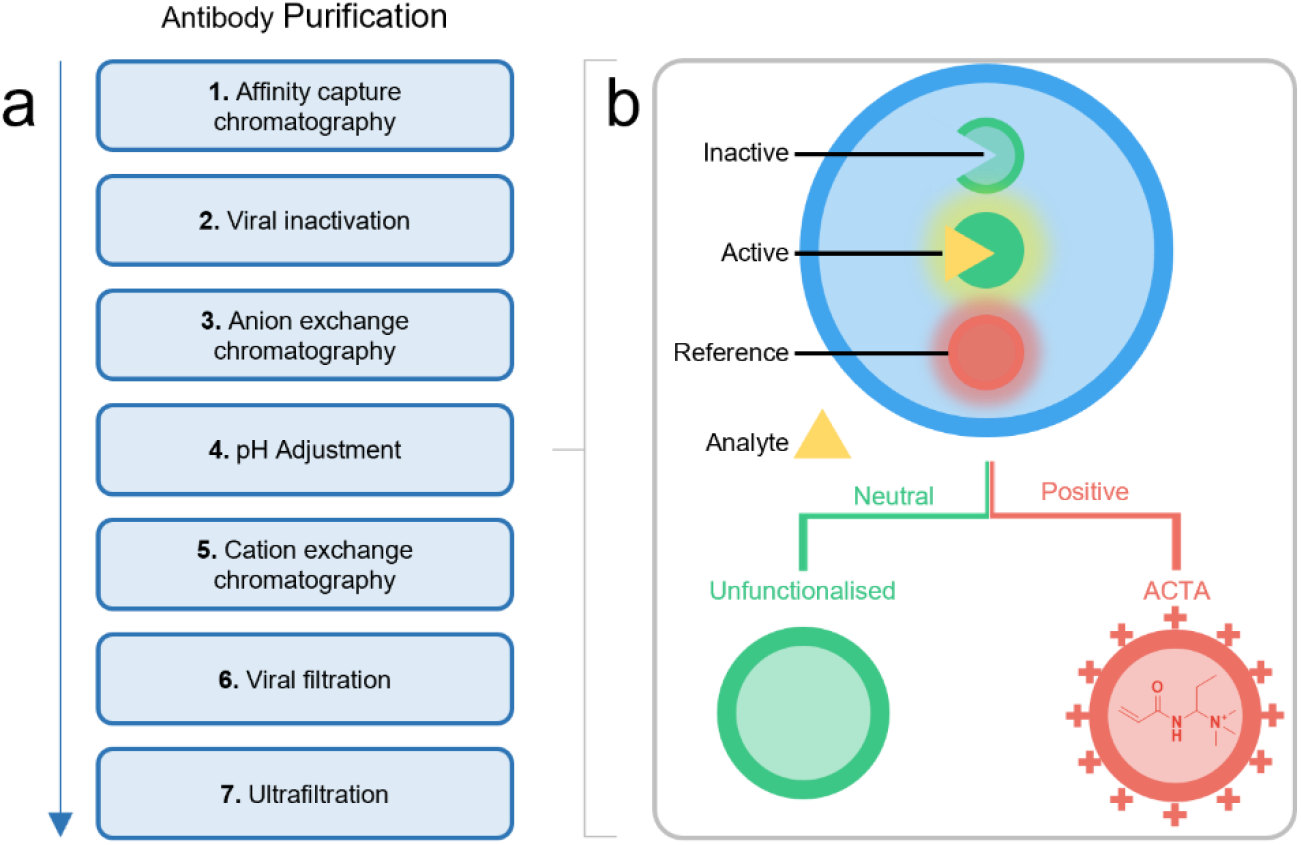
Antibody purifying workflow and schematic of pH-sensitive nanosensors. **a** Workflow of a typical process for purifying monoclonal antibodies, which could benefit from pH control during low volume downstream optimisations. **b** Schematic diagram of fluorescent nanosensors and how they can be functionalised with acrylamidopropyltrimethyl ammonium hydrochloride (ACTA) to produce nanoparticles with a positive zeta potential.

Fluorescent nanosensors are inert, versatile biosensors, that can be used to make measurements of key molecules and ions in microenvironments of interest, such as low volume downstream processing, and are ideal for real-time measurements of dynamic processes [9]. Due to the 1) combination of analyte responsive and reference fluorophores, 2) their small biocompatible matrix (∼50 nm diameter), and 3) application of the highly analyte-sensitive principles of fluorescence, nanosensors demonstrate accurate ratiometric analyte quantification as well as high spatial and temporal measurements (**Fig. 1b**, [10])

We have developed nanosensors to quantify a variety of analytes which include pH [11] molecular oxygen [12] and temperature [13] in complex model systems. The pH-sensitive fluorescent nanosensors have gathered the greatest momentum, due to their ability to perform accurate ratiometric measurements, using a combination of pH-sensitive and pH-insensitive fluorophores that are bound to inert polyacrylamide matrix (**Fig. 1b**). This approach allows for precise pH measurements through standard fluorometric detection. The matrix is also chemically versatile and can incorporate molecules that can alter their surface charge properties, permitting differential interaction with their microenvironments. For example, acrylamidopropyltrimethyl ammonium hydrochloride (ACTA), can be used to confer a positive zeta potential to the surface of unfunctionalised particles, which are typically neutral in nature [14].

Fluorescent pH sensitive nanosensors have been evaluated and validated in a range of complex and diverse microenvironments and demonstrated their immense potential by 1) mapping the acidification in nematode model organisms (*Caenorhabditis elegans* [15] & *Pristionchus pacificus* [16]), 2) elucidation of subcellular fermentation pathways in *Saccharomyces cerevisiae* [14], 3) determining the intracellular processing of foreign material in human mesenchymal stem cells (hMSCs) [17] and 4) characterising the evolution of acid by-products during bacterial biofilm growth [18]. Thus, fluorescent nanosensors, potentially provide a solution to HTPD for bioprocessing.

This study evaluates the application of pH-sensitive nanosensors for HTPD of downstream experimental conditions using a TECAN™ automated liquid handling platform. The application of fluorescent nanosensors will be explored through sacrificial sampling, by analysing discrete samples from the process stream. These studies are anticipated to address the critical need in scale-down process development for efficient, accurate, automated, low-volume and high-throughput monitoring of pH. By harnessing the capabilities of pH-sensitive nanosensors and integrating them with automated platforms, this study seeks to boost efficiencies in downstream bioprocessing, particularly in antibody purification.

## MATERIALS AND METHODS

### MATERIALS

#### Nanosensors

Acrylamide 99% minimum, N,N’methylenebis(acrylamide) and Brij L4, ammonium persulfate (APS), dioctyl sodium sulfosuccinate salt (AOT), and N,N,N’,N’-tetramethylethylenediamine (TEMED) were purchased from Sigma-Aldrich (Gillingham, United Kingdom). Hexane HPLC grade and Ethanol analytical grade were obtained from Fisher Scientific (Loughborough, United Kingdom). Argon gas acquired from BOC Gases (Manchester, United Kingdom). Deionised water (18.2 MΩ) was generated by Elga Purelab Ultra (ULXXXGEM2).

#### Fluorophores

5-(and-6)-carboxyfluorescein succinimidyl ester (FAM-SE) Oregon Green 488 carboxylic acid succinimidyl ester (OG-SE), and 5-(and-6)-carboxytetramethylrhodamine succinimidyl ester (TAMRA-SE) were purchased from Invitrogen™ (Paisley, United Kingdom). N-(3-aminopropyl)methacrylamide hydrochloride (APMA) was obtained from Polysciences Inc (Warrington, United Kingdom). Sodium borate decahydrate was obtained from Sigma-Aldrich (Gillingham, United Kingdom).

#### pH buffer solutions

Sodium phosphate dibasic and citric acid monohydrate were purchased from Sigma-Aldrich (Gillingham, United Kingdom).

#### Chromatography & Antibodies

a monoclonal antibody, (Poros™ HS 50, Thermofisher) and strong anion exchanger (Capto™ Q, Cytiva) resins.

## METHODS

### Preparation of fluorophores to N-(3-aminopropyl) methacrylamide

APMA (5 mg, 0.028 mmol) was dissolved in a sodium borate tetradecahydrate buffer solution (2.5 mL, 50 mM, pH 9.5). Aliquots the APMA stock solution (0.200 mL, 0.002 mmol) were added to the fluorophores FAM-SE (0.001 g, 0.002 mmol) OG-SE (0.001 g, 0.002 mmol) and TAMRA-SE (0.001 g, 0.002 mmol) in separate vials. The vials were stirred continually overnight to provide fluorophores sufficient time to conjugate to APMA (FAM-APMA, OG-APMA & TAMRA-APMA). Aliquots of conjugate stock solutions were used for nanosensor synthesis.

### Extended dynamic range pH-sensitive nanosensor synthesis

Surfactants Brij 30 (3.080 g, 8.508 mmol), dioctyl sodium sulfosuccinate salt (1.590 g, 3.577 mmol) and deoxygenated hexane (42 mL) were stirred under argon for 10 min. FAM-APMA (15 μL, 5 mg/mL), OG-APMA (15 μL, 5 mg/mL), TAMRA-APMA (60 μL, 5 mg/mL), acrylamide (0.540 g, 7.58 mmol), and *N,N*′methylenebis(acrylamide) (0.160 g, 1.307 mmol) were dissolved in deionised water made up to 2 mL. For positively charged particles acrylamide (0.513 g, 7.22 mmol), *N,N*′methylenebis(acrylamide) (0.152 g, 0.986 mmol) and acrylamidopropyltrimethyl ammonium hydrochloride (ACTA, 75% w/v 119 μL, 0.401 mmol) were dissolved in deionised water made up to 2 mL. These monomer solutions were added to the stirring hexane surfactant solution and allowed to deoxygenate for a further 10 min. Polymerization initiators ammonium persulfate (30 μL, 10% w/v) and *N,N,N*′,*N*′-tetramethylethylenediamine (15 μL, 0.1 mmol) were added to the stirring solution to initiate polymerization. The mixture was left to stir for 2 h under argon, after which hexane was removed *via* rotary evaporation. Nanoparticles were precipitated and washed with ethanol (30 mL) using centrifugation (10 times, 6000 rpm, 10 min), with a Hermle centrifuge (Z300). After the final wash, the pellet was resuspended in a small amount of ethanol (10 mL) and rotary evaporated until dry. Nanoparticles were stored in a light protected container at 4 °C.

### Dynamic light scattering (DLS)

DLS was performed using a Malvern Zetasizer™ Nano ZS (Malvern Panalytical, Royston). The system is equipped with a 5 mW He-Ne laser source (633 nm), operating at an angle of 173°. Fluorescent pH-sensitive nanoparticles were suspended in deionised water (1 mg/mL). Measurements (n=3, 25 °C) were made using a disposable polystyrene cuvette (Sarstedt, Leicester). The mean hydrodynamic diameter of the particles was computed from the intensity of the scattered light using Malvern Zetasizer software (6.12).

### Zeta potential

Nanoparticles were suspended (1 mg/mL) in filtered phosphate buffered saline solution (1 in 10, NaCl 13.7 mM KCl 0.27 mM Na_2_HPO_4_ 1 mM KH_2_PO_4_ 0.18 mM, 0.02 μm, Millipore). Nanoparticle suspensions were transferred to zetasizer cuvettes (DTS1061, Malvern), flushed with filtered deionised water. Zeta potential measurements were made in triplicate (constants used to make measurements of polyacrylamide nanoparticles: refractive index: 1.452, parameters used for dispersant deionised water dispersant; refractive index: 1.330, viscosity: 0.8872 cP, dielectric constant: 78.5 εr, Model Smoluchowski F (Ka) 1.5). All samples were allowed to equilibrate to 25 °C for 120 s prior to measurement.

### Calibration

The calibration process involved preparing a stock solution with a concentration of 2 mg/mL nanosensor in deionised water, which was then diluted with sodium phosphate dibasic and citric acid monohydrate buffer solutions. pH changes of buffer solutions to temperature were evaluated, which was monitored against measurement performed in plate reader (**Table S1**). Calibration buffers and nanosensors were mixed in a 1:1 ratio and the max concentration achievable in the absence of detector saturation was monitored (**Fig. S1**). Final concentrations of 1 mg/mL and 10 mg/mL nanosensor were selected for measuring pH in 100 μL and 25 μL volumes, respectively, which provided sufficient signal in the absence of detector saturation. Nanosensors both with neutral and positive surface charge measured the Green Intensity (OG/FAM excitation 470 nm, emission 520 nm) / Red Intensity (TAMRA excitation 540 nm and emission 577 nm) ratio. A comparative analysis evaluated the sensitivity and the measurement accuracy between the two sensor types (**Table 1 & Fig. S2**)

**Table 1.**
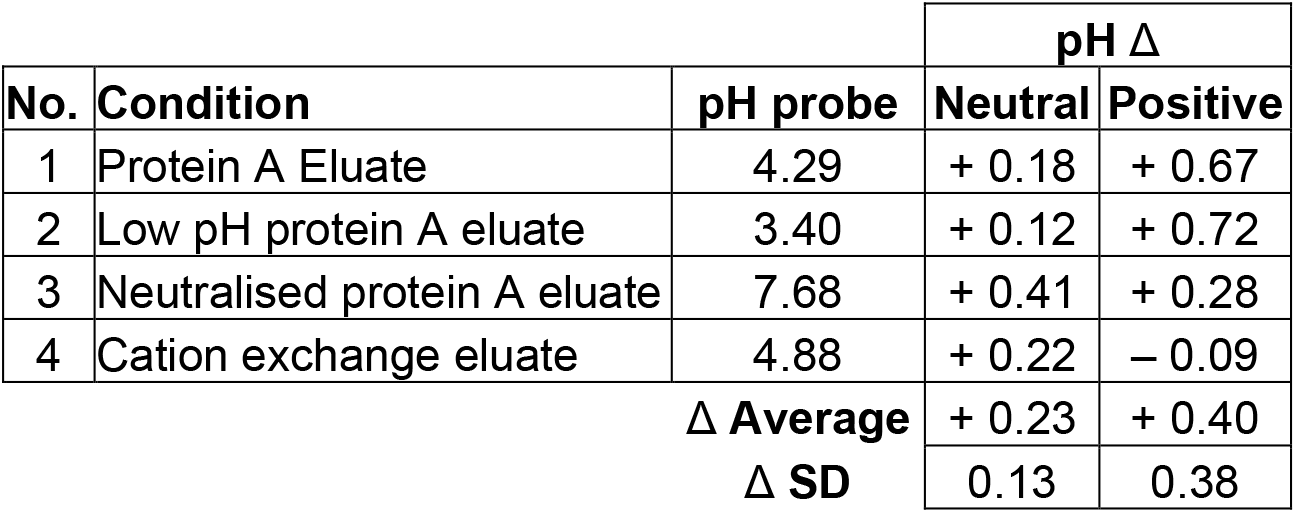
Protein A Purification Column Validation. Comparison of eluate fractions collected from subsequent monoclonal antibody purification stages. pH probe measured vs fluorescent nanosensor pH measurement, calculated by converting measured fluorescence signal ratios to pH based on calibration data obtained using calibration solutions with known pH. Deviation measured as pH delta from probe measurement (n=3).

### Fluorescence measurement and validation of pH

Sample preparation was performed using a TECAN Freedom EVO^®^ 200 liquid handling system controlled by an EVOware^®^ 2.8 SP3 software. Fluorescence measurement was performed in 12.5-100 μL sample volumes collected in 384-well plates using a TECAN Infinite^®^ M1000 Pro microplate reader at the corresponding wavelengths for each fluorophore. Automated dispensing of low-volume pH indicator solutions is highlighted in **Sup. Video 1**. Fluorescence intensity data was captured using Magellan™ software V 7.2 by TECAN™. The background fluorescence of the samples was measured without any nanosensors. The pH of the calibration buffers and the intermediate samples were validated using a Mettler Toledo Seven Compact pH meter.

### Intermediate sample preparation

A bispecific monoclonal antibody was expressed extracellularly in a Chinese hamster ovary (CHO) cell line generated at AstraZeneca, Gaithersburg, MD, USA. Cells were separated from the conditioned medium using depth filtration and the clarified harvest was purified using Protein A chromatography (MabSelect Sure™) to generate the protein A eluate intermediate sample. The protein A eluate was first titrated to pH 3.40 to generate the low pH protein A intermediate and subsequently titrated to pH 7.68 to produce the neutralised protein A intermediate. Following that anion exchange (Capto™ Q) and cation exchange (Poros™ 50 HS) chromatography was performed sequentially, to generate the cation exchange chromatography eluate. The pH of process intermediate samples of the bispecific monoclonal antibody was evaluated using nanosensors. 12.5-50 μL protein A chromatography eluate, low pH protein A eluate, neutralised protein A eluate and cation exchange eluate in sodium acetate-based buffer were diluted with each of the nanosensor neutral and positive stock solutions (2 mg/mL, 1:1 ratio) to a sample volume of 25-100 μL (final nanosensor concentration 1 mg/mL).

### Characterisation of nanosensor adsorption to chromatography resins

Nanosensor binding to anion exchanger (Capto™ Q, Cytiva) and cation exchanger (Poros™ 50 HS, Thermofisher) was investigated using batch binding experiments. A constant resin volume of 50 μL of each resin was first equilibrated with 50 mM sodium acetate, pH 5.0 for cation exchange resin and 50 mM Tris, pH 7.0 for anion exchange resin. Following the removal of the equilibration buffer, the resins were incubated for at least 30 min with nanosensors added at 0.1 mg/mL concentration and protein aliquots loaded at increasing concentrations by varying the added volume from 100 μL to 500 μL in increments of 100 μL. The resin-protein-nanosensor suspension was first washed with their corresponding equilibration buffer and eluted with 50 mM Tris, 500 mM NaCl, pH 7.0 for anion exchange resin and 50 mM sodium acetate, 500 mM NaCl, pH 5.0 for strong cation exchange resin. Fluorescence intensity was measured in the effluent for the Capto™ Q sample, in the effluent collected during the product loading and salt strip phase, and for the Poros™ 50 HS sample, collected during the binding and elution phase. The measured fluorescence intensity was corrected for background fluorescence using resins equilibrated in their corresponding equilibration buffers in the absence of nanosensors.

## RESULTS & DISCUSSION

### Nanosensor characterisation

To investigate size and charge of synthesised nanoparticles, DLS and zeta potential analysis was conducted. DLS particle analysis revealed that neutral and positively functionalised nanoparticles had hydrodynamic diameters centred at 32.67 nm and 37.84 nm, respectively, with size distributions ranging from 15.69 nm to 105.70 nm (**Fig. 2a, Table S2**). Comparative analysis of nanoparticle surface charge indicated significantly different zeta potentials for neutral (−5.08 ± 4.05 mV) and positive nanoparticles (+12.87 ± 1.25 mV, P<0.001, Student’s t-test, **Fig. 2b**).

**Fig. 2.**
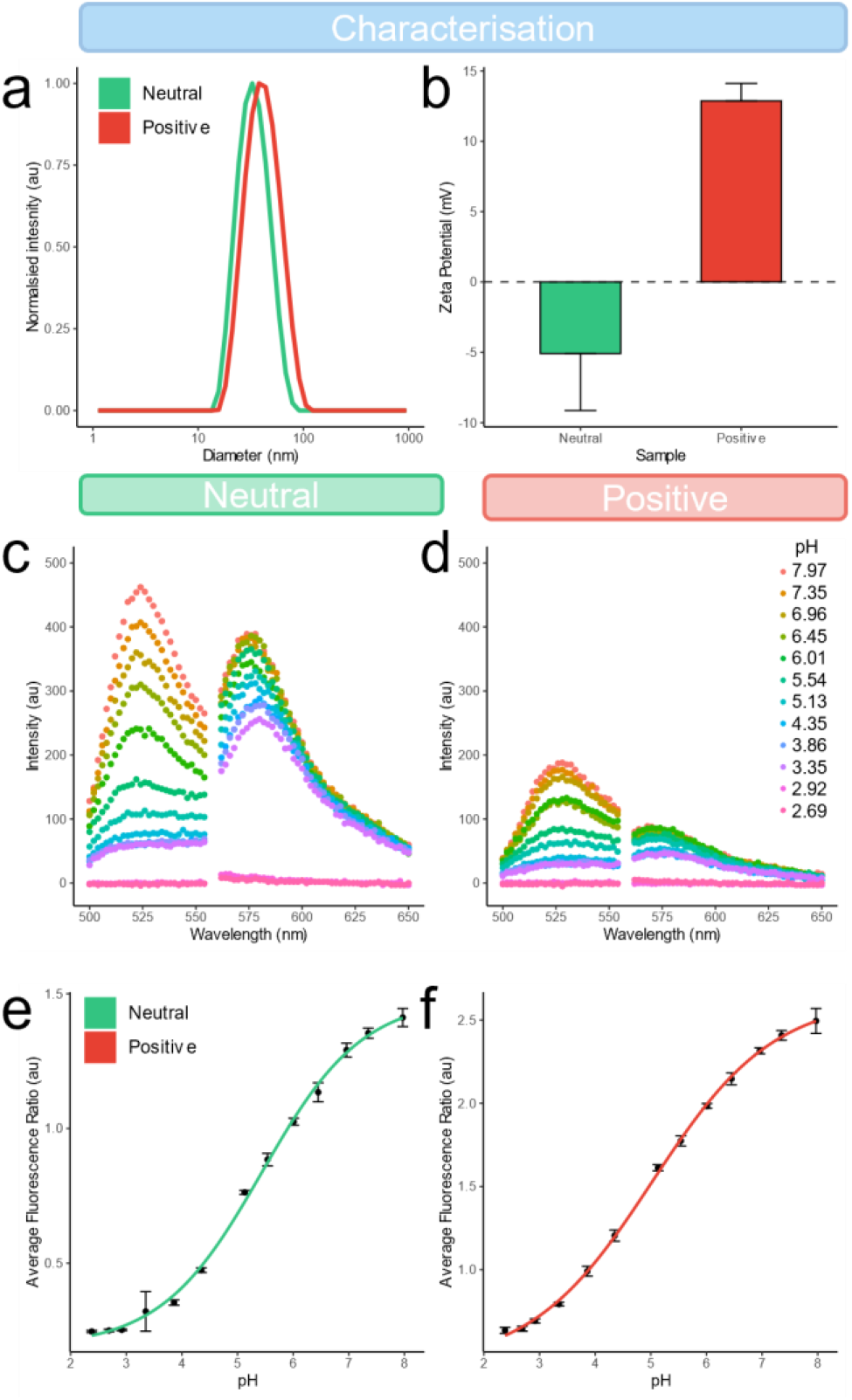
Characterisation of fluorescent pH-sensitive nanosensors. Comparative **a** DLS intensity size distributions and **b** zeta potentials for fluorescent nanosensors (1 mg/mL) suspended in PBS (1:10 dilution, pH 7.4, n=3). Fluorescence spectra (**c** neutral, **d** positive), calibration curves (**e** neutral, **f** positive), for fluorescent nanosensors (100 μl, 1mg/mL, 384-well plate, error refers to standard deviation, n=3).

To evaluate the pH-dependent fluorescence response of neutral and positively charged nanosensors, spectroscopic analysis was conducted. Both types of nanosensors, neutral (**Fig. 2c**) and positive (**Fig. 2d**), demonstrated increased fluorescence emission with pH when using analyte-sensitive fluorophores: OG (for pH < 5.5) and FAM (for pH > 5.5). In contrast, the reference fluorophore TAMRA showed relative insensitivity to pH changes, aligning with previous studies [12]. Interestingly, positively functionalised nanosensors exhibited a 2.75x and 4.44x reduction in green and red emission intensities, respectively. This reduction was attributed to the addition ACTA functional group, which competes for binding on the nanoparticle matrix with APMA-functionalised fluorophores. This leads to fewer fluorophores binding to the nanosensor matrix and a decrease in emission signal. Calibration curves (neutral **Fig. 2e**, positive **Fig. 2f**), were produced by extracting averaged fluorescence intensity at each measured pH value and fitted with a four-parameter regression equation [11]. Despite variations in fluorescence intensity, the calibration curves for neutral and positive sensors remained consistent, as confirmed by normalised calibration curves (*P*<0.001, **Fig. S2**). This consistency suggests that the ratio of OG & FAM to TAMRA fluorophores remained constant, despite changes in total observed emission intensity.

### Nanoparticle resin binding

To assess whether the differential surface zeta potentials of neutral and positively functionalised nanosensors would affect their binding and release to cationic and anionic chromatography resins, the nanoparticle retention was investigated using process buffers typically used for monoclonal antibody purification. Ion exchange chromatography separates molecules based on their net surface charge and is dependent on surrounding pH. Cation exchange chromatography operates in “bind-and-elute” mode, where the positively charged region of the target molecule will bind to the negatively charged ligands carried by resin through electrostatic interactions. Poros™ 50 HS, a strong cation exchange resin, is functionalised with sulfonate groups (-SO_3_^-^), enabling it to electrostatically attract and bind positively charged molecules. Whereas Capto™ Q, a strong anion exchange resin with a quaternary ammonium group (-N^+^(CH_3_)_3_), the quaternary ammonium group binds negatively charged species on to its positively charged ligand. This makes Capto™ Q suitable for operating in “flowthrough” mode, capturing negatively charged species, such as DNA, endotoxins, while letting the target molecule flow through the column with the mobile phase.

In our experiments, we collected samples from two stages of the chromatography process – 1) binding/flowthrough and 2) elution/salt-strip phases. During the binding phase, a substantial proportion of neutral particles were detected in the flowthrough for both Poros™ 50 HS (66 ± 15%, **Fig. 3a**) and Capto™ Q (98 ± 27%, **Fig. 3b**). In contrast, positively charged particles were predominantly adsorbed by Poros™ 50 HS, with only a small proportion remaining unbound (4 ± 5%, **Fig. 3a**), the positively charged particles are recovered in the elution/salt-strip phase. The majority of both neutral and positive nanosensors were observed only in the flowthrough for the Capto™ Q (97 ± 31%, **Fig. 3b**). These results suggest that positively charged nanosensors have the potential to be used as an online, in-process pH detection tool that can be added to the bioprocess at the beginning and removed after the last chromatography step, enabling pH control in an automated, high throughput manner. This differential binding efficiency of nanosensors to Poros 50™ HS and Capto™ Q resins, therefore, highlights their specificity and potential for targeted applications in biomolecule separation and purification.

**Fig. 3.**
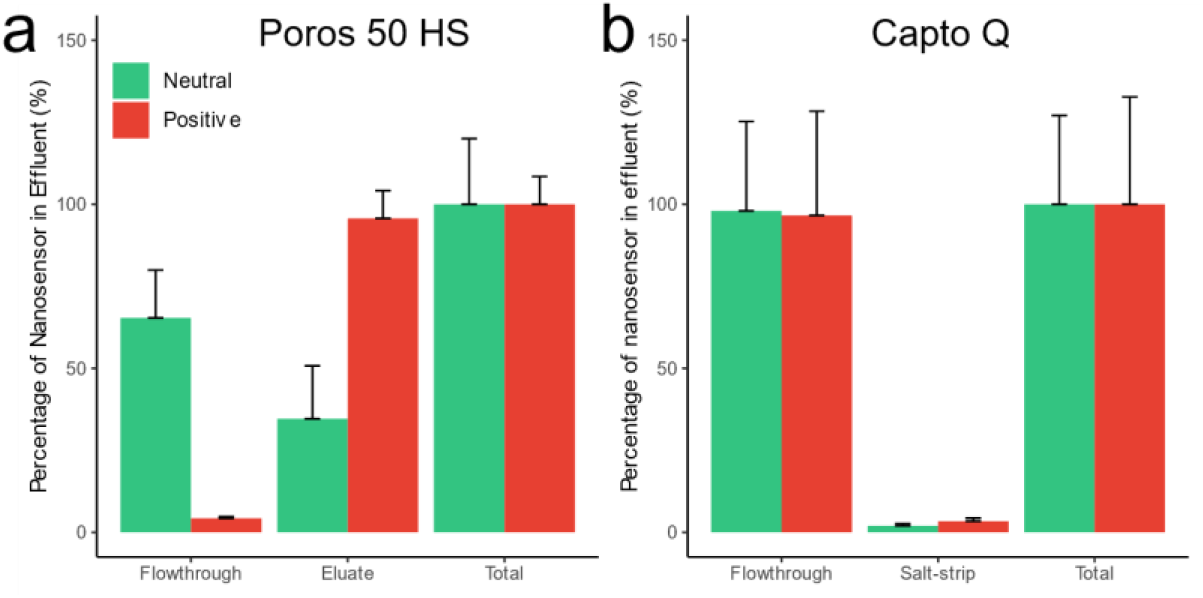
Differential nanoparticle resin binding. Comparison of fraction of neutral and positive fluorescent nanosensors released from **a** Poros 50 HS cation and **b** Capto Q anion exchange resin. Data normalised to total fluorescent nanosensor recovered (error = SD, n=3)

### Downstream Process Automation

Building upon these findings, an automated scale-down pH adjustment workflow was developed, using a TECAN™ liquid handling platform (**Fig. 4**), which allows for high-throughput and precise measurements of pH changes during the purification stages. This workflow, designed for microtiter plate-based pH evaluation, leverages the capabilities of nanosensors for assessing protein A purification process performance involving low-volume (12.5 μL) samples of bispecific monoclonal antibodies (**Fig. 5**).

**Fig. 4.**
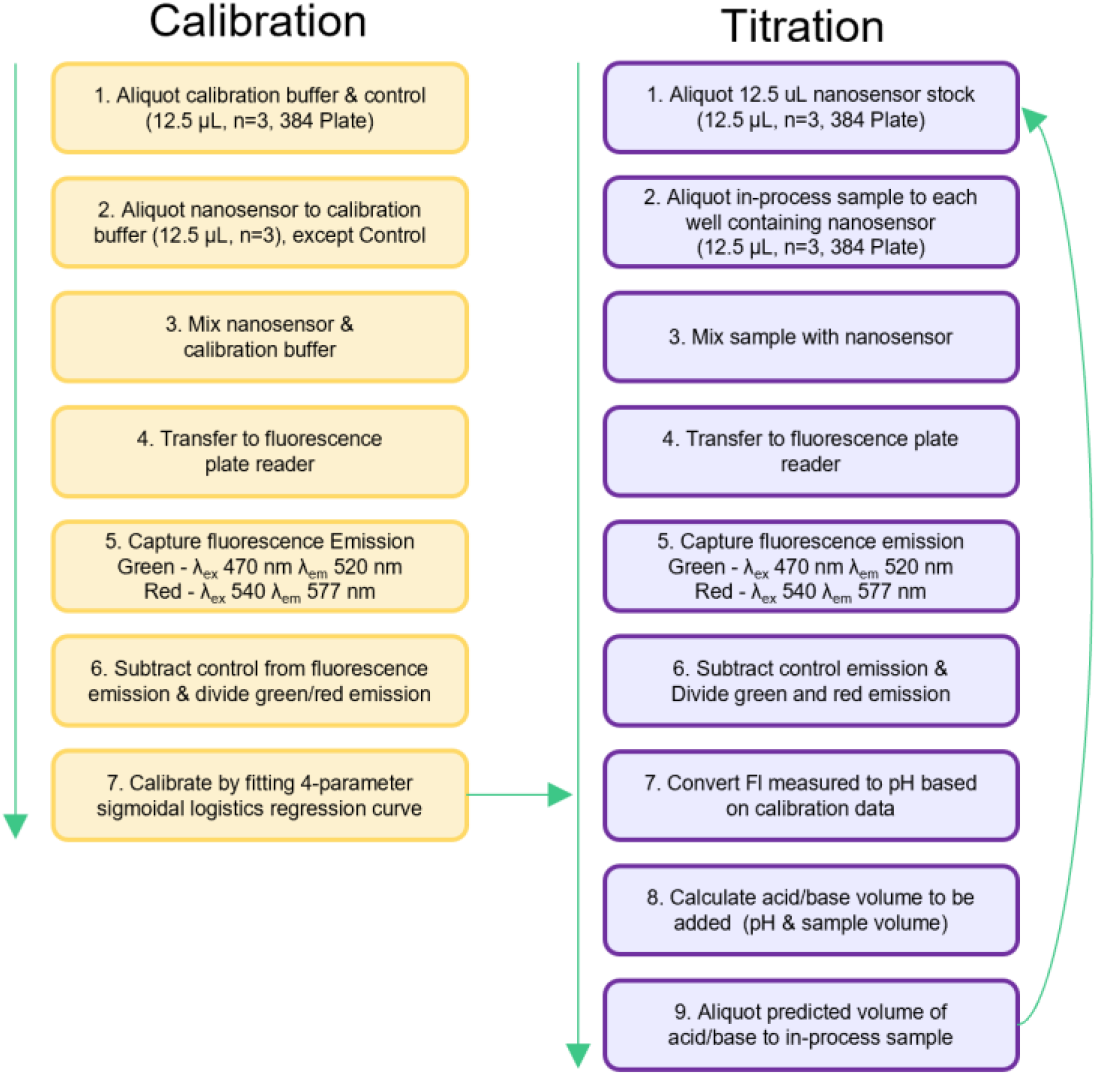
Automation optimisation. Workflow for the calibration and optimisation of the sacrificed measurement of pH using automated protein purification pathways and fluorescent nanosensors.

**Fig. 5.**
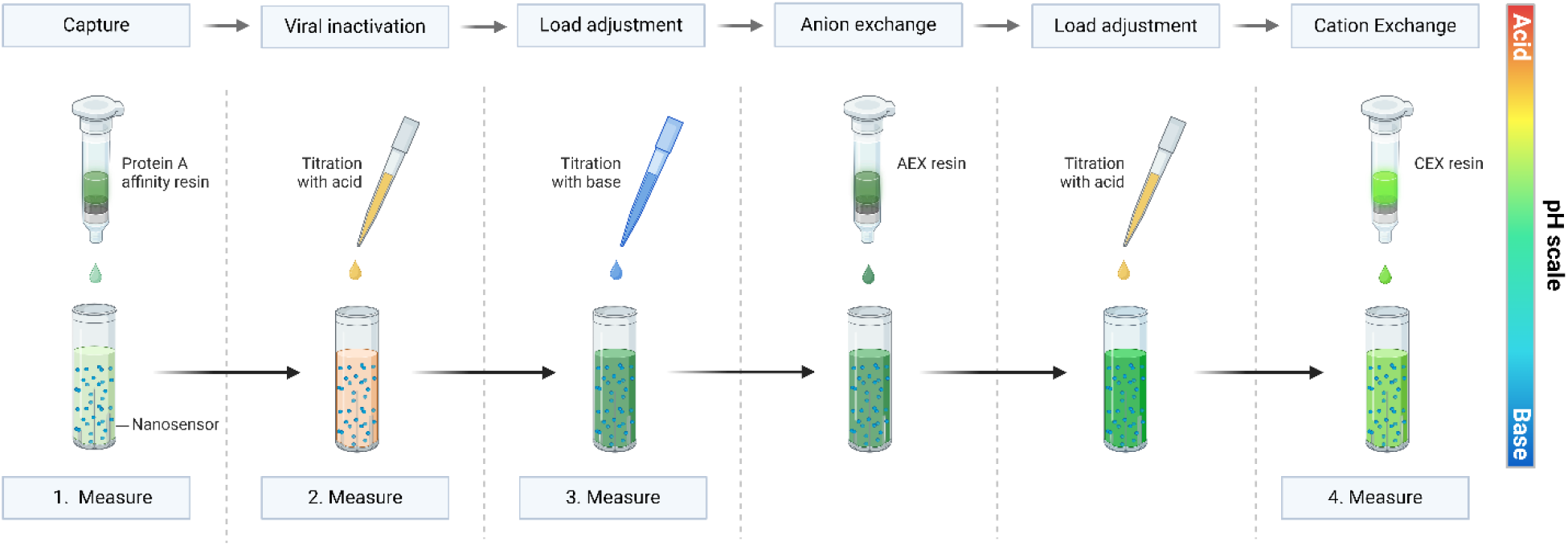
Automated TECAN workflow for antibody purification using nanosensors. Microwell plate-based, high throughput pH measurement of protein purification process intermediates in 25 μL sample volumes. The nanosensor particles suspended in the sample allow for fast, accurate determination of the pH using a fluorescence spectrophotometer.

In evaluating the pH of intermediate samples from the monoclonal antibody purification pathway using nanosensors (**Table 1**), neutral fluorescent nanosensors were able to effectively characterise pH when compared to traditional pH probe measurements. They exhibited an average deviation (Δ) of +0.23 ± 0.13 across all stages. The most significant deviation occurred for the evaluation of post protein A eluate neutralisation, with a Δ of +0.41, likely due to the pH measurements exceeding the nanosensors dynamic range (pH > 7.5). Whereas positive nanosensors exhibited in general higher average deviation in measured pH across the protein purification process (Δ of +0.40 ± 0.38). This larger deviation with positive nanosensors could be attributed to their lower relative emission compared to neutral sensors, potentially leading to a greater noise-to-signal error and, consequently, larger deviations. We also observed a general data trend, where nanosensors generally overestimated pH values compared to probe-based methods. This observation was true for all data except for the positively charged nanosensors at the cation exchange stage, where a decrease of pH -0.09 Δ was recorded. Importantly, yield determination of the protein A capture step is not influenced by nanosensor presence when absorbance was measured at 280 nm (**Fig. S3**). Therefore, the technological advance we have demonstrated here represents a significant step forward in streamlining HTPD sample analysis for continuous monitoring of pH.

## CONCLUSION

In conclusion, this study has successfully demonstrated the capabilities and applications of fluorescent nanosensors in pH monitoring and adjustment during automated HTPD. Neutral and positively charged nanosensors were produced, which demonstrated differential binding behaviour with ion exchange resins. This may permit at line measurements, as the nanosensors could be present in the product stream for continual pH analysis and be separated at a specific stage without affecting the final product quality. Fluorescent nanosensors demonstrated effective pH measurement, when compared to a pH probe, using automated low-volume sacrificial sampling at each stage of the protein purification process. This has not been possible before for small-scale HTPD, due to the lack of compatible suitable, and accurate measurement technologies. Future research will focus on optimising these nanosensors for broader applications and addressing the challenges observed in the dynamic range and signal-to-noise ratios, especially in the context of positively charged nanosensors. Our study highlights the promising role of nanosensors in enhancing the efficiency and accuracy of scale-down HTPD and builds a solid foundation for the development of improved monitoring of biopharmaceutical manufacturing processes.

## Author Contributions

Conceptualization: GCB, AT, JWA, VMC

Methodology: GCBC, AT, VMC

Investigation: GCB, AT, VMC

Visualization: VMC

Supervision: AT, JWL, VMC

Writing: – original draft: GCB, AT, VMC

Writing: – review and editing GCB, AT, JWA, VMC

## ADDITIONAL INFORMATION

### Supplementary Information

is available for this paper, which includes Supporting Information PDF, and Video.

### Correspondence and requests for materials

should be addressed to VMC.

