## Supporting Information for "Fluorescent pH-sensitive nanosensors enable precise low-volume monitoring in high-throughput bioprocess manufacturing"

Supporting Figures S1-3
Supporting Table S1-2
Caption for Supporting Video 1

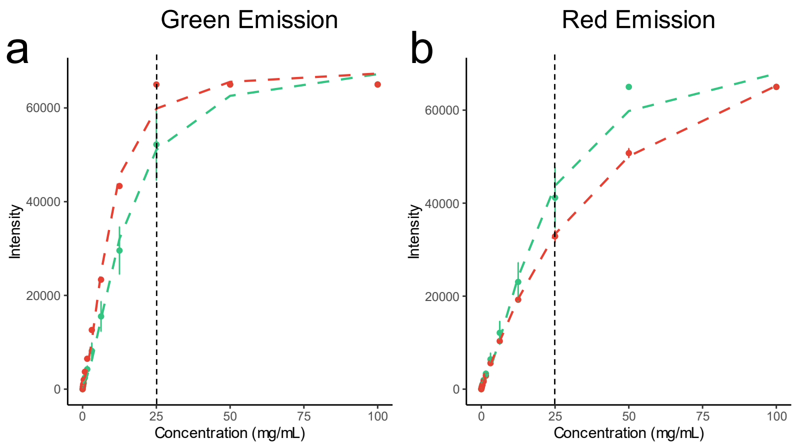


**Fig. S1 | Nanosensor concentration vs emission intensity.** Comparative **a** green and **b** red emission intensities for fluorescent nanosensors at concentrations ranging from 0.1 to 100 mg/mL at pH 7.2. Max nanosensor concentration identified as 25 mg/mL in 96 well plate avoiding signal saturation.

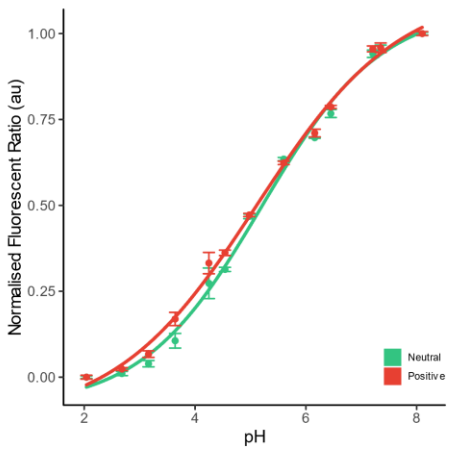


**Fig. S2 | Nanosensors normalised calibration curves.** Comparative normalised neutral and positive pH calibration curves.

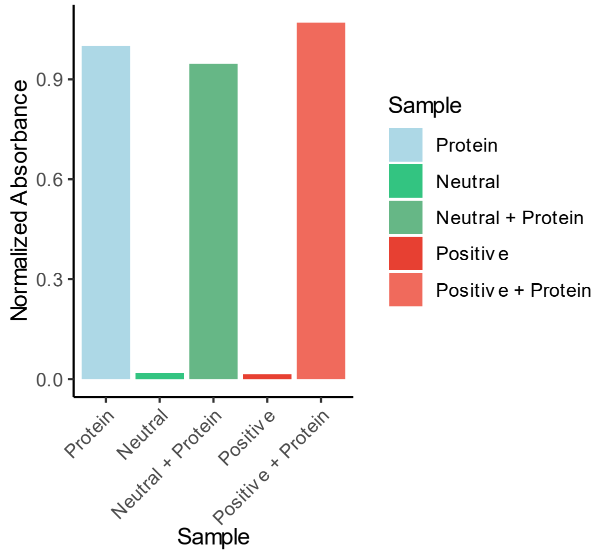


**Fig. S3 | Protein and Nanosensor absorbance.** Comparative normalised absorbance of protein in the presence and absence of neutral and positive nanosensors (1mg/mL, A280).

**Table S1 | Temperature dependent pH**. Characterizing the pH differences from pH meter at 21.7 °C and 19.6 °C.

| | **pH @ 21.7** | **pH @ 19.6** | **Change** |
|---|---|---|---|
| **1** | 7.97 | 7.90 | -0.07 |
| **2** | 7.35 | 7.27 | -0.08 |
| **3** | 6.96 | 6.88 | -0.08 |
| **4** | 6.45 | 6.35 | -0.10 |
| **5** | 6.01 | 5.91 | -0.10 |
| **6** | 5.54 | 5.42 | -0.12 |
| **7** | 5.13 | 5.04 | -0.09 |
| **8** | 4.35 | 4.28 | -0.07 |
| **9** | 3.86 | 3.81 | -0.05 |
| **10** | 3.35 | 3.29 | -0.06 |
| **11** | 2.92 | 2.86 | -0.06 |
| **12** | 2.69 | 2.58 | -0.11 |
| **13** | 2.38 | 2.18 | -0.20 |

**Table S2 | Sizing studies.** Dynamic Light Scattering hydrodynamic average diameters, standard deviation and polydispersity index (PDI) (n=3).

| | **Distribution Centres (nm)** | **Average diameter (nm)** | **Standard Deviation** | **PDI** |
|---|---|---|---|---|
| **Neutral** | 32.67 | 31.42 | 0.18 | 0.102 |
| **Positive** | 37.84 | 38.37 | 0.14 | 0.116 |

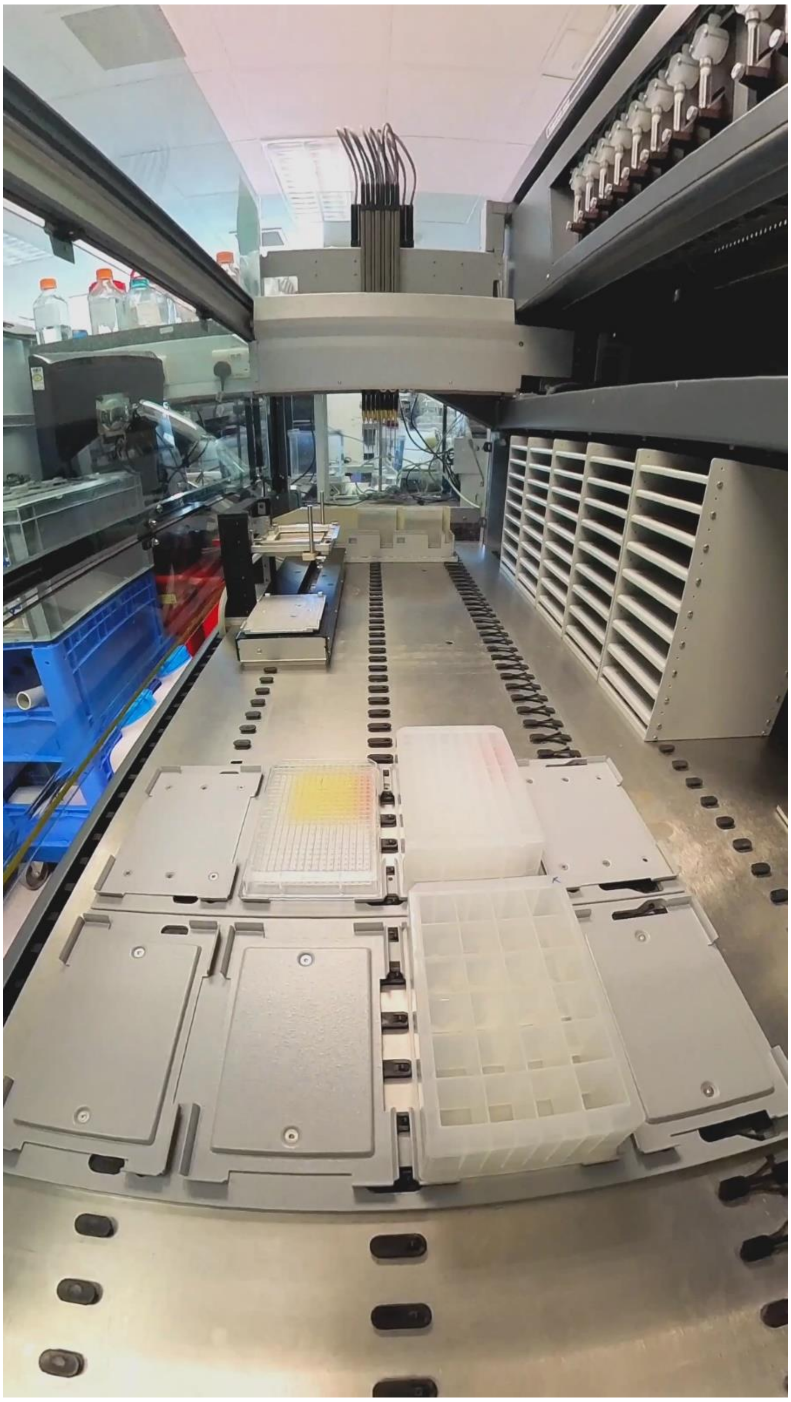

**Supplementary Movie 1.** Automated dispensing of low-volume pH indicator solutions using the TECAN Freedom EVO® 200 liquid handling platform.
